# Head-Centred Sound Localisation Behaviour Reveals Head and World Centered Reference Frames in Auditory Cortex

**DOI:** 10.64898/2025.11.29.691317

**Authors:** Nadia Aghili, Stephen Town, Jennifer Bizley

## Abstract

The auditory system relies on head-centered localisation cues to estimate sound source position. Yet listeners and single neurons can encode sound in world-centred spatial reference frames. However, most studies of sound localisation constrain the subject’s orientation within a fixed speaker ring, making it difficult to distinguish which reference frame underlies spatial coding.

We utilised a sound localisation task that required ferrets localise sound location, relative to the head, across rotations in the starting platform within the testing arena. Two ferrets were trained to discriminate front from back sounds. Once trained, the orientation of both the central platform and the target speakers were rotated between testing sessions, retaining the task contingency in head-centered coordinates, but breaking the alignment of head and world centered reference frames. Probe trials were presented from the other speakers so that spatial receptive fields could be constructed and compared across central platform rotations.

Neural activity was recorded from auditory cortex using chronically implanted 32-channel arrays. Across 1,777 recorded units, 596 were stimulus driven. Spatially tuned neurons were predominantly head-centred, with 208 and 232 units classified as head-centred in the two animals, compared with 48 and 49 world-centred units and 33 and 26 un-tuned units, respectively. Temporal analysis showed that most units retained the same reference-frame classification from onset to offset, but the magnitude of their head-versus-world preference diminished over time, with offset responses shifting towards more ambiguous and, in some cases, world-centred coding.

Together, these results demonstrate that auditory cortex maintains parallel head centered and allocentric representations that are head dominated at onset but incorporate increased world-centred information at offset, supporting perceptual constancy during dynamic listening.

## 1. Introduction

Accurate sound localisation is fundamental for survival, communication, and navigation. Yet, because the head and body are continually in motion, the auditory system must generate representations of sound location that remain stable across movement. This problem is often framed in terms of spatial reference frames, where sound location may be encoded relative to the listener (egocentric, head-centred) or relative to the external environment (allocentric, world-centred) ^1^. Distinguishing between the two frames is difficult, and this is mirrored in broader work on spatial cognition. Imagined translations, which rely on egocentric bearings, are updated accurately, whereas imagined rotations, which require allocentric updating of orientation, tend to produce larger errors and slower responses, especially when real movement cues are absent ^2^.

In the auditory system specifically, sound position is initially coded in head-centred coordinates ^3^, consistent with an egocentric frame. However, behavioural and developmental evidence indicates that stable interaction with the environment also requires allocentric coding, as shown by preserved localisation performance across postural changes in adults^4^ and by the gradual emergence of this ability in children ^5^. These observations raise the question of how reference frames identified in behaviour are instantiated in the brain.

Extensive neurophysiological work has characterized how sound localisation cues are extracted in the midbrain ^6,7^ and transmitted to higher-order regions including auditory cortex^8,9^, parietal cortex, and prefrontal cortex^10^. Among these regions, auditory cortex is pivotal for accurate localisation in primates and carnivores ^11^ but the coordinate frame of spatial representation under naturalistic conditions remains unclear, despite the recognised computational and perceptual value of allocentric coding ^12–14^ . Studies in freely moving animals have addressed this directly, revealing that although many auditory cortical neurons encode sound location in head-centered coordinates, a subset encode it relative to stable positions in the external environment, providing direct evidence for world centered coding ^15, 16^ .

These findings suggest that world centered coding can emerge in auditory cortex under specific behavioural conditions. Building on this work, our previous behavioural study ^16^ introduced a head-centred task in which ferrets discriminated broadband noise bursts presented from in front of versus behind the head, while the rotation of the central platform was systematically varied across sessions to manipulate the relationship between head- and world-centred coordinates and assess the contribution of world-centred information to the task.

Here, we extend this work by analysing neuronal activity recorded while animals performed the task. By recording across multiple platform rotations, we were able to classify neurons as head-centred, world-centred, or ambiguous (mixed). Consistent with our findings in passively listening animals, we observed a predominance of head-centred units, alongside a minority of neurons that represented sounds in world-centred coordinates or exhibited mixed reference frames.

## 2. Method

### 2.1 Experimental Procedures

The experimental setup has been described in detail previously; here we provide a concise summary, with full descriptions of the behavioural task available in^16^.

#### 2.1.1 Ethics statement

All experimental procedures were approved by local ethical review committees (Animal Welfare and Ethical Review Board) at University College London and The Royal Veterinary College, University of London and performed under license from the United Kingdom Home Office (Project Licenses 70/7267 and PPPP1253968) and in accordance with the Animals (Scientific Procedures) Act 1986.

#### 2.1.2 Animals

Two pigmented female ferrets (0.5-3 years old, 600-1100 g) were housed in groups under enriched conditions. Animals were trained to report sound location for water rewards under a regulated water schedule, with daily monitoring of body weight and demeanor alongside fluid intake to ensure welfare. Regular otoscopic examinations confirmed ear health.

#### 2.1.3 Experimental design

Experiments were conducted in a double-walled, sound-attenuating chamber (IAC Acoustics; 1.6 × 1.6 × 1.1 m) lined with 45-mm acoustic foam and dimly illuminated with white LED strip lights. Within the chamber, animals were tested in a custom-built circular arena (0.5-m radius) enclosed with plastic mesh. A ring of twelve loudspeakers (Visaton FRS8) was positioned at 30° intervals, 55 cm from the arena centre (Fig. 1). At the arena centre, a central platform provided a controlled starting position. The platform contained three lick ports (left, centre, right) for trial initiation and responses. When the animal occupied the central port, its head was aligned with the centre of the loudspeaker array. For the head-centred task used here, the platform was rotated between sessions to sample multiple locations (up to 12), in 30° increments, around the arena centre (Fig. 1a), thereby dissociating the orientation of the animal from the fixed positions of the loudspeakers within the arena. This design ensured that sounds were always delivered from the trained speaker locations corresponding to fixed directions relative to the head (e.g., front vs back, shown in Fig. 1a in blue and red, respectively), while their positions in the external world (Fig. 1b) varied systematically with platform angle. Speakers were calibrated and matched in level (give B&K specs). Occasionally we performed physical swaps of pairs of speakers to demonstrate that animals were performing the task based on sound location and not based upon characteristics of the speakers themselves.

**Figure 1.**
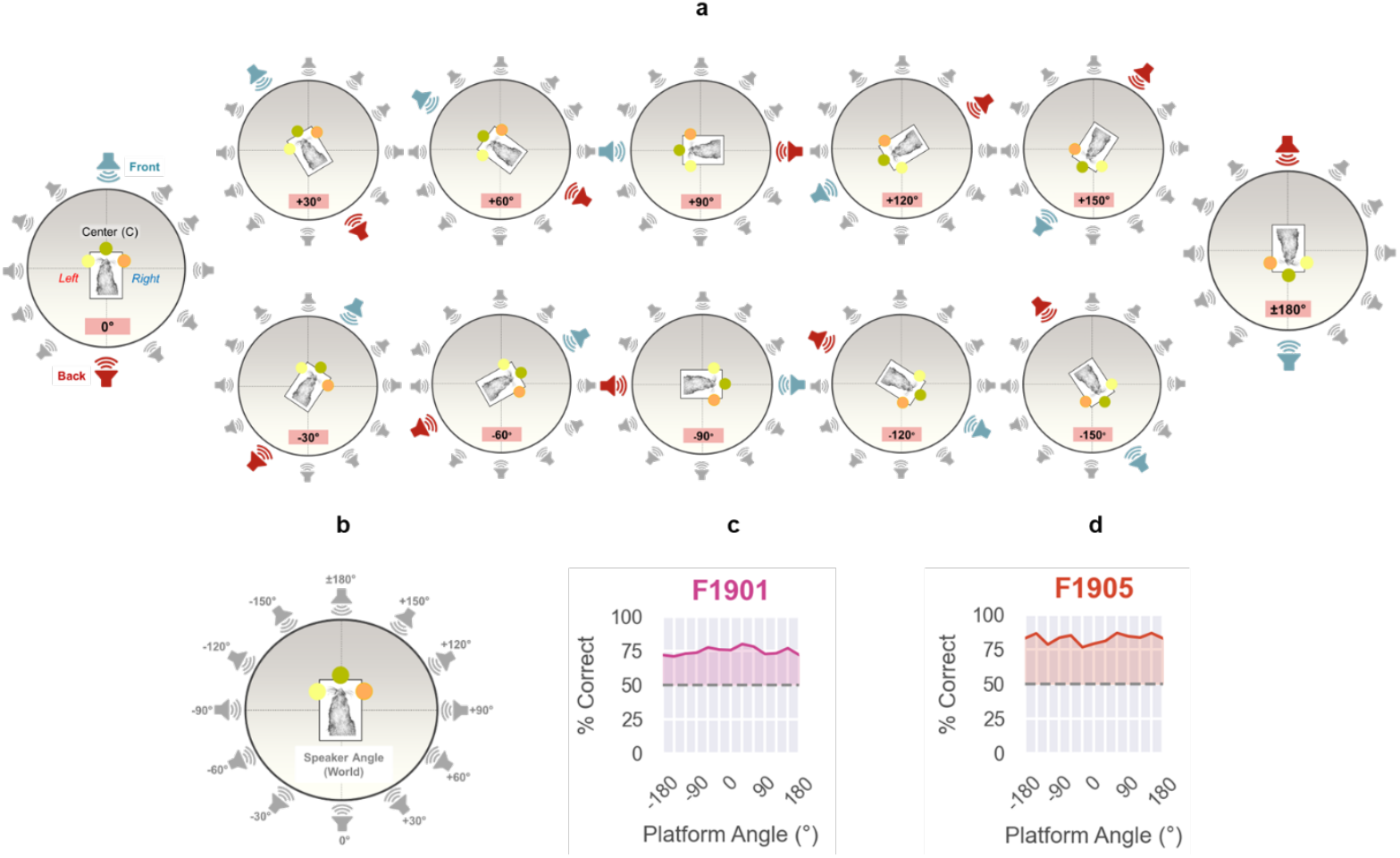
Experimental set-up and behavioural performance. (a) Schematic of the behavioural arena showing the ferret at the central spout while the platform is rotated to different orientations. Grey speakers indicate probe locations spaced at 30° intervals, and the ferret initiates the task on the central rectangular platform; the blue speaker marks the front target and the red speaker the back target. (b) Speaker angles defined relative to the arena (world-centred) coordinate system. (c-d) Behavioural performance (% correct responses) for the two ferrets across different platform orientations.

**Figure 2.**
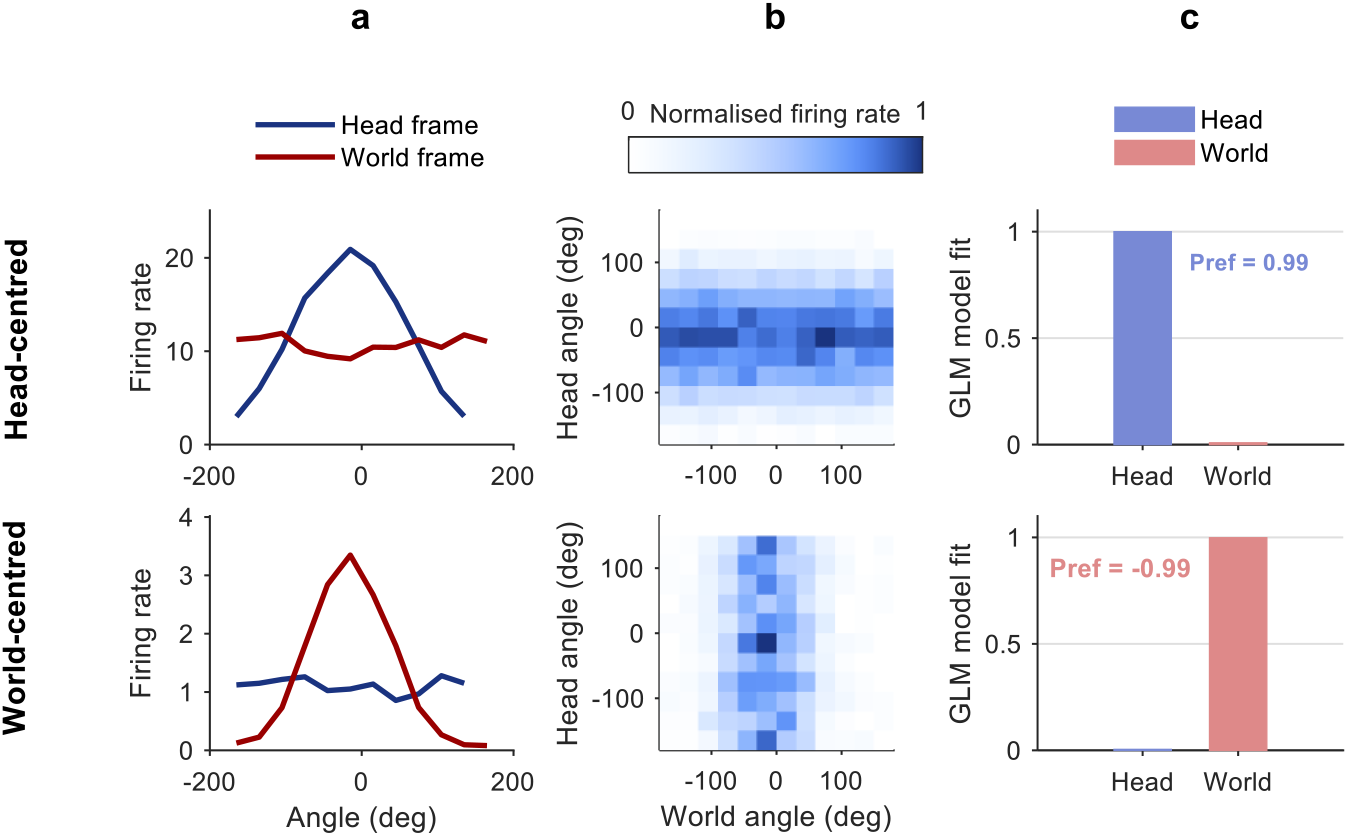
Simulated head (top) and world (bottom) centered coding across coordinate frames. (a) Simulated spatial receptive fields in head and world coordinates. (b) Head by world heatmaps of mean firing rate for each unit type. (c) Quantitative model comparison, where bars show Model Fit for the head angle (by purple) and world angle (by pink) GLM and the numbers (Pref) report Model Preference as defined in Equation 5.

#### 2.1.4 Stimuli

On each trial, animals-initiated presentation of a single broadband noise burst (250 ms duration, 60-63 dB SPL) generated afresh on each trial. On a subset of trials (≤10%), probe sounds were presented from untrained loudspeaker positions, shown in grey in Fig. 1a (i.e. grey positions other than the trained front-back axis), and responses to these were always rewarded. Continuous low-level broadband noise (45 dB SPL) was played through all speakers to mask extraneous sounds.

#### 2.1.5 Head-centered localisation task

##### Task design

Ferrets were trained to discriminate sounds based on their position relative to the head midline. Two animals (F1901 and F1905) were trained to distinguish sounds originating directly in front (180°) from those directly behind (0°). Their behavioural performance exceeded 70% correct responses across different platform angles for both ferrets, as shown in Fig. 1c-d. To ensure stable head position and minimize dynamic localisation cues, ferrets were required to hold at the central lick port for 0.2-0.7 s before stimulus onset; trials were only initiated if the animal remained in position throughout the sound. Trial initiation triggered the presentation of a 250-ms broadband noise burst (60-63 dB SPL) was presented. Responses were made at one of two lateral lick ports on the platform, positioned 60° to the left and right of the centre platform (shown as yellow and orange circles in Fig. 1a at platform orientation 0°): front sounds were rewarded with a right-port response, and back sounds with a left-port response.

##### Training

Training proceeded in stages. Initially, animals received repeating noise bursts from left or right of the head while holding at the central port, and responses at either lateral port were rewarded. Once animals reliably initiated and responded, error contingencies were introduced so that left sounds required Left responses and right sounds required Right responses. Across sessions, this sound-response mapping was rotated counterclockwise until animals discriminated front (Right response) versus back (Left response). The task was then made progressively more demanding by reducing stimuli to a single 250-ms burst, increasing the required hold time before stimulus onset, and extending timeouts following errors. Animals typically required 2-3 months to reach stable performance (>70% correct) on the single-burst task.

##### Testing

After training, the central platform was rotated to vary the orientation of the ferret relative to the loudspeaker array. Rotations were introduced gradually, beginning with small angles (30°) and extending to a full 180°. Animals were then tested across the full 360° range of platform orientations in 30° increments, with rotations pseudorandomly interleaved across sessions. Within a session animals completed many trials with a fixed platform orientation (typically 50-300).

#### 2.1.6 Neuronal recording

Animals were implanted bilaterally over primary and secondary auditory cortex with 32 channel WARP-32 electrode drives (Neuralynx) loaded with high impedance tungsten electrodes (FHC). After surgery electrodes were individually lowered into auditory cortex. Electrodes were further advanced into the brain every few weeks so that across 1-2 years of recordings, sampling was obtained from all cortical layers in each ferret, with final depth reached once electrodes extended beyond the estimated lower border of auditory cortex. Neural activity in auditory cortex (both hemispheres) was recorded continuously during behavioural testing using chronically implanted 32-channel wireless WARP devices. Voltage signals were recorded and digitised using a MultiChannel Systems 2100 wireless recording device with a sample rate of f 20 kHz. On average, data was recorded across 8.8 depths (range: 7 to 10), with 17.5 sessions per depth (range: 15.5 to 21).

### 2.2 Simulation

Alongside the real recordings, we generated simulated neurons with spatial receptive fields designed to express head centered or world centered tuning; in both cases, spike probability was modelled as a function of stimulus location

#### Head centered

Head centered tuning described the relationship between spike probability (*P*) and sound source angle relative to the subject’s head midline (*θ*). Spike probability was modelled in MATLAB (MathWorks) using a Gaussian function:

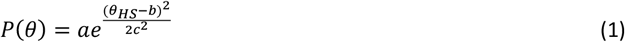

Where *a* is the peak amplitude, *b* the preferred angle (set to 0° at the head midline), and *c* the tuning width. In our case, parameters (*a* = 1.044, *b* = 0, *c* = 75.7) were determined by manual fitting to obtain qualitatively matched head centered and world centered tuning functions.

##### World centered

World centered tuning described the relationship between spike probability (*P*) and sound source position in world-centred Cartesian coordinates (*x, y*). Spike probability was simulated as the dot product of two logistic probability functions defined separately for the x- and y-axes:

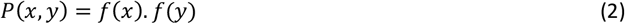

With

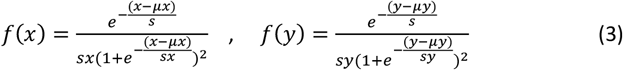

Parameters were set to *ux* = 1000, *sx* = 400, *uy* = 0, *sy* = 1000. Sound sources were arranged on a circle of radius 1000 units, with Cartesian coordinates computed from their polar positions in the world.

##### Simulation details

In both models, 2000 trials were simulated. On each trial, head direction was randomly sampled from discrete orientations at 30° increments between -180° and 180°, and sound sources were sampled from loudspeakers positioned every 30° around the circular array (radius 1000). This ensured that the same world-fixed source could correspond to different head-centred angles across trials, replicating the behavioural paradigm (Section 2.1.3). Spike probability functions were linearly rescaled to firing rates between 0 and 20 spikes/s. Poisson spike counts were generated with means equal to these firing rates. For visualisation, firing rate maps were constructed by averaging responses within 30° bins of head and world angle, and rates were normalised within session to their maximum.

### 2.3 Data analysis

Spike sorting was performed using Python (version 3.11), and all post-sorting analyses were conducted in MATLAB (version R2024b).

#### 2.3.1 Spike sorting

Recordings were organized into group of blocks (where each block was a single testing session), with each group corresponding to the same electrode depth within a single hemisphere. In the spike-sorting pipeline, the raw voltage trace from each block was first evaluated for data quality by calculating its signal power. The median power across all blocks was used as a reference and blocks whose power fell outside the range between zero and twice the median was excluded from subsequent analyses. Then, each retained blocks contained 32-channel raw voltage traces, which were processed sequentially through a standardized preprocessing pipeline.

First, channels with abnormally high power were excluded; specifically, channels whose mean absolute power exceeded three times the median across all channels were identified as noisy and removed. These typically corresponded to damaged electrodes. Next, transient peaks occurring immediately after prolonged static periods, arising from temporary disconnection of the battery or recording headstage, were corrected. Flat segments lasting more than 0.4 s (in addition to ±20-sample padding) were replaced with the local median amplitude to suppress sharp step edges that could otherwise be misclassified as spikes. To detect these residual events, we then applied a chunk-based approach in which the mean amplitude within each chunk was compared to the channel median; chunks whose mean exceeded twice this threshold were replaced with the median of the recording. Finaly, the cleaned signals were then re-referenced using a common average reference to reduce shared noise, band-pass filtered in the spike band (300-3000 Hz), and whitened to equalize noise variance across channels.

Preprocessed sessions recorded at the same groups of depth were concatenated to form continuous datasets, which were then fed to spike sorting using MountainSort4 (MS4) ^17^ via the SpikeInterface framework ^18^ . Key parameters included an adjacency radius of 20, with filtering and whitening disabled to avoid redundant processing, and 10 worker threads for parallelization. Throughout this study we use the term ‘units’ to refer to the resulting sorted (but likely multi-unit) clusters.

#### 2.3.2 Spatial tuning features

Spike clusters identified by MS4 were evaluated for sound responsiveness using a Poisson generalized linear model (GLM)^19^ . For each unit and recording session, spike counts were measured within four temporal windows relative to stimulus onset: a baseline period (-50 to 0 ms), an early response window (0 to 50 ms), a late pre-offset window (200 to 250 ms), and a late post-stimulus window (250 to 300 ms). Two comparisons were performed: baseline versus early, and late pre-offset versus late post-stimulus. For each comparison, two GLMs were fitted; a null model assuming a constant firing rate across time windows, and a full model including an epoch term indicating whether each spike count occurred before or after stimulus onset (baseline=0, post=1 as predictor). Improvement in model fit was assessed with a likelihood-ratio test, and responses were considered significant when the full model provided a better fit than the null model (*p* ≤ 0.05). To ensure stability and exclude weak effects, analyses were limited to sessions with at least 20 trials, and only changes exceeding a standardized effect size of |dz| ≥ 0.2 (Cohen’s dz) were accepted.

Units were classified as stimulus-driven if they exhibited significant modulation in at least two recording sessions within each group of depth. Finally, a post-selection power filter was applied to remove units with very weak responses. For each unit, we computed the median peak firing rate across GLM-passing sessions from smoothed peri-stimulus time histograms, and units whose median peak rate was less than 20% of the across-unit median were excluded. The resulting dataset comprised reliable, stimulus-driven units suitable for further analysis.

A subset of stimulus-driven neurons (units) whose responses were systematically modulated by sound location, expressed in either head-centered or world-centered coordinates, were classified as spatially tuned, as assessed with generalized linear mixed-effects models (GLMMs) ^20^. Trials from all sessions were pooled, and for each trial, head-centred and world-centred sound angles were calculated based on speaker location and the animal’s head rotation. Spike counts were measured within two post-stimulus windows (0-150 ms and 250-400 ms after onset). Angles were represented as sine and cosine predictors to capture circular tuning. For each unit, Poisson GLMMs were fitted, each including a random intercept for session, in order to account for session-to-session changes in baseline firing rates. In the first instance two models were constructed; firstly, a model with a constant term only (null model) and secondly a full more that had both head- and world-angle as predictors. A likelihood-ratio test determined whether the full model provided a significantly better fit (*p* ≤ 0.05), indicating spatial tuning.

For visualisation of reference-frame tuning, we generated reference-frame heatmaps for each unit. To account for session-wise differences in baseline firing rate, trials were first partitioned by platform orientation (session). For each rotation, we constructed a head by world firing-rate matrix (30° bins, 0-0.25 s), normalised it to the range 0-1, and then combined the per-rotation maps by the element-wise maximum to obtain a single aggregate heatmap per unit, with world angle on the x axis and head angle on the y axis.

#### 2.3.4 Analysis of head-centre and world-centre Units

For each unit that showed significant spatial tuning, two reduced Poisson GLMMs were fitted containing either head- or world-centred predictors, each with a random intercept for session. The model with the lower Akaike Information Criterion (AIC) was taken as the unit’s preferred reference frame and used for categorical classification as *head*- or *world*-centred.

To further quantify tuning strength and population-level biases, we calculated continuous model-fit metrics derived from GLMM deviance, following our previous approach ^1^. Four models were compared: a constant (null) model, a head-centred model, a world-centred model, and a full model that included both head and worlds angles as predictor (each with session fit as a random effect). For each coordinate frame (test), model fit was defined as the deviance improvement of the corresponding model over the constant model, normalized by the improvement achieved by the full model:

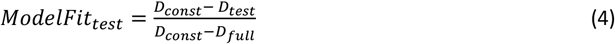

The deviance of the constant (null) model is denoted as *D*_*const*_, that of the full model as *D*_*full*_ and *D*_*test*_ refers to the deviance of either the head-centred or world-centred model. Model fitting was performed separately for early (0-150 ms) and late (250-400 ms) post-stimulus windows. When units showed significant tuning in both periods, the early window was used for classification and model-fit analysis; if tuning was absent early but significant later, the late window was used instead.

We then calculated the model preference as:

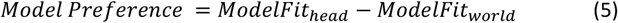

Model preference values ranged from -1 (indicating a better fit to sound angle in the world) to +1 (indicating a better fit to sound angle relative to the head). We also applied a threshold of |0.2| to the model-preference index to indicate strongly head- or world-centred units, a convention used only for visualisation and shown later in Fig. 4; categorical classifications themselves were determined solely by AIC according ^1^.

**Figure 3.**
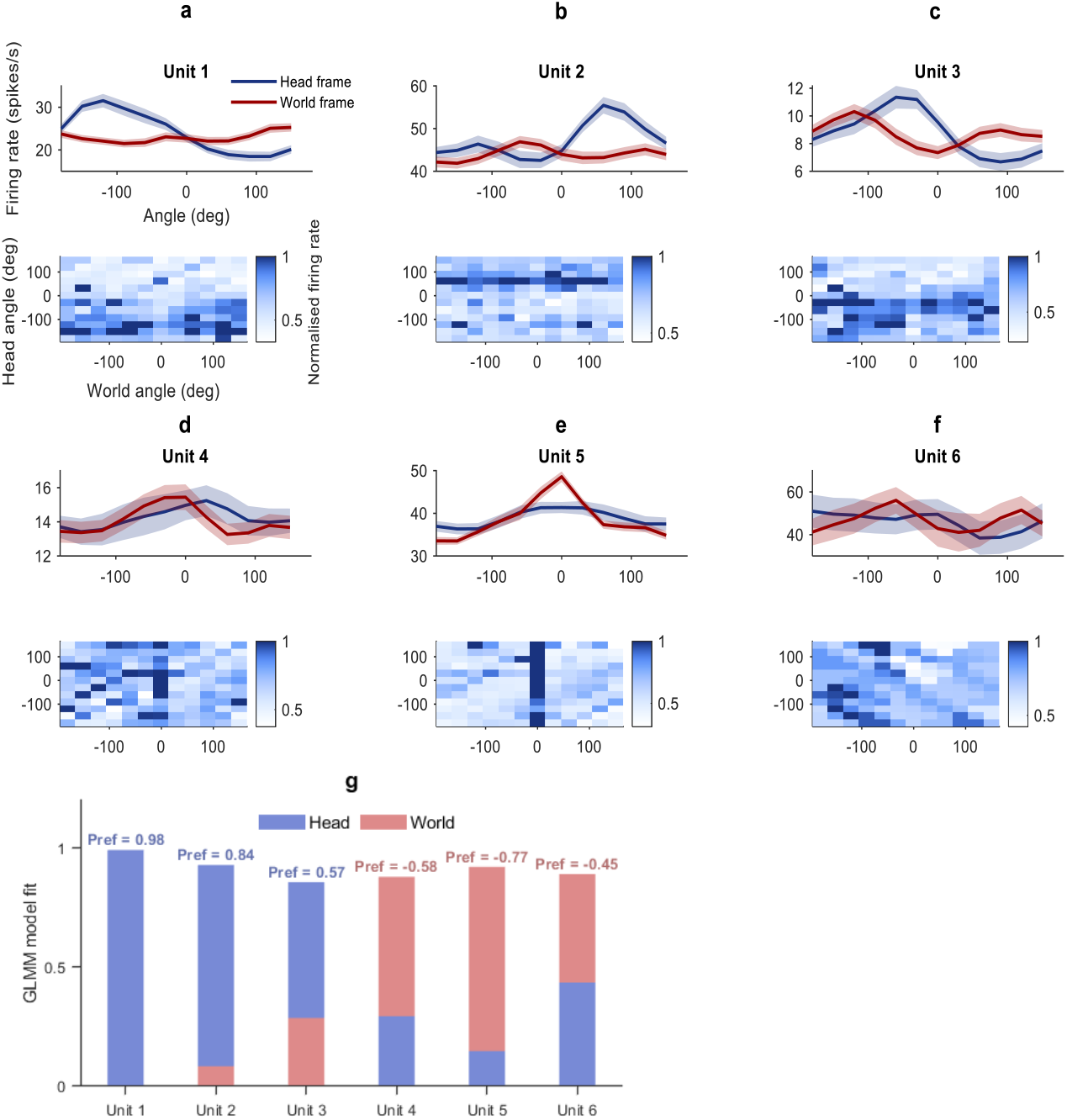
Reference frames in neural recordings. (a) Spatial receptive fields in head-centred and world-centred coordinates, and head-by-world firing-rate maps, for six units recorded in an example ferret (F1905). The navy line shows the head-centred frame and the dark red line the world-centred frame; shaded regions indicate ± SEM. Units 1-3 show head-centred structure, units 4-5 show world-centred structure, and Unit 6 shows mixed tuning. (b) GLMM fits for head angle (purple) and world angle (pink). Numbers above each pair of bars give the model-preference value for each unit, confirming the assigned reference frame.

**Figure 4.**
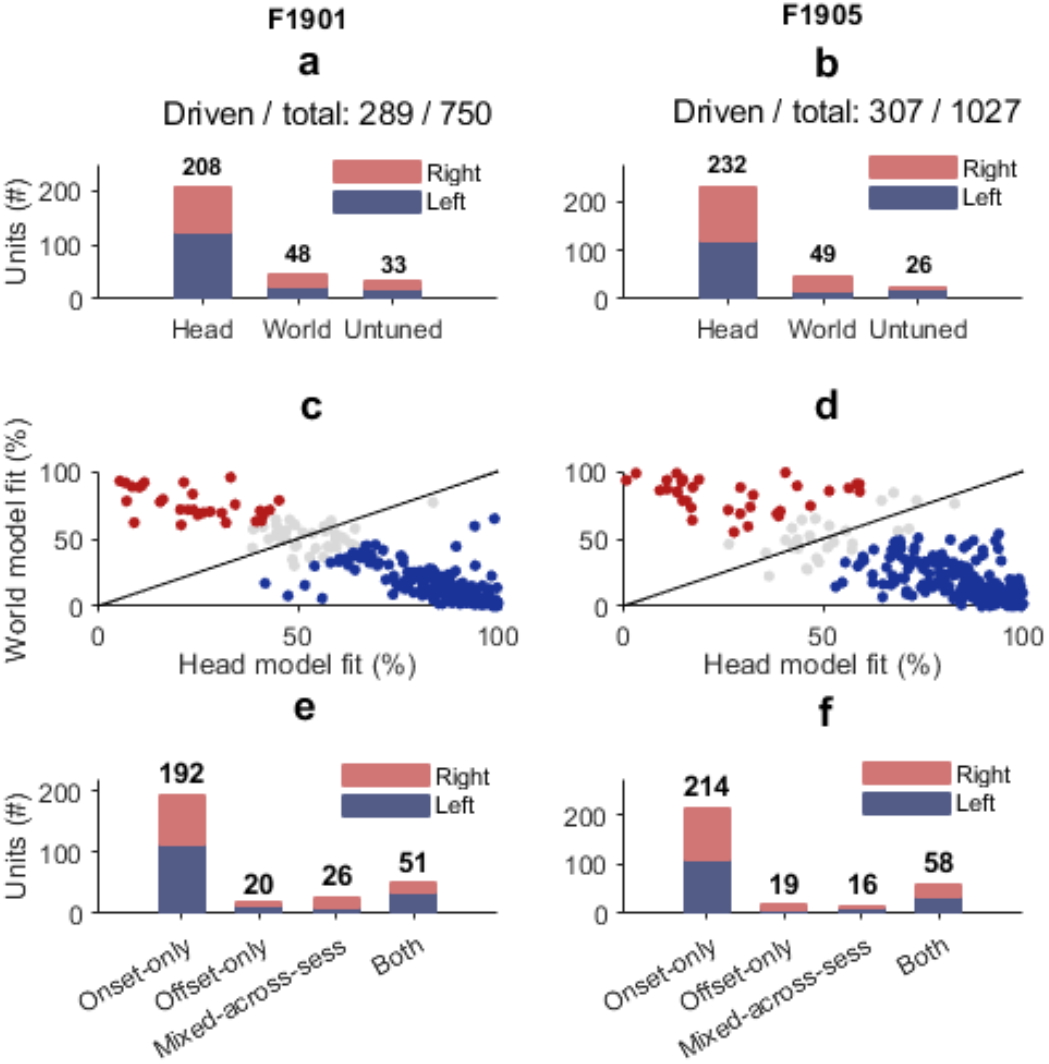
Summary of population reference-frame. (a, b) Population tuning in the early window. For F1901 (a), stimulus-driven and total units (left–right–total) are 166/413, 123/337 and 289/750, respectively. For F1905 (b), stimulus-driven and total units are 156/439, 151/588 and 307/1,027. Bar heights indicate the number of head-centred, world-centred and not-tuned units; navy bars show left hemisphere and pink bars show right hemisphere. (c, d) Model-fit comparison for tuned units in the early window. For each ferret (F1901 in c, F1905 in d), scatter plots show model fit (%) for the head model (x-axis) versus the world model (y-axis); the diagonal line indicates equal fit. Points are coloured according to model preference: navy, preference > +0.2 (strong head-centred); pink, preference < -0.2 (strong world-centred); grey, |preference| ≤ 0.2 (ambiguous). (e, f) Temporal categories of stimulus-driven units (both early and late window). For F1901 (e) and F1905 (f), bars show the number of units classified as onset-only, offset-only, mixed across sessions or both. Within each bar, navy and pink indicate left and right hemispheres, respectively.

#### 2.3.5 Temporal Comparison of Onset and Offset Spatial Responses

Units were first screened using the 50 ms session-wise responsiveness tests described in Section 2.3.2. Only units that showed significant modulation in both onset and offset phases, and met the across-session criterion, were included in the temporal comparison. For each included unit, spatial tuning over a 150 ms window was then assessed separately for onset and offset, pooling trials across sessions and fitting Poisson GLMMs with a random intercept for session (Section 2.3.3). Neurons were first gated with a likelihood-ratio test comparing the Full and Constant models (α = 0.05). Neurons that passed this gate were classified as head or world according to the lower AIC and labelled mixed (ambiguous) when the model-preference index was smaller than |0.2|; neurons that did not pass the gate were labelled as untuned. Within-unit agreement between onset and offset was then quantified by comparing the time-specific labels, without introducing any criteria beyond those defined in Sections 2.3.2 and 2.3.4.

## 3. Results

Animals were trained to localise sounds in a head-centered reference frame (Figure 1a) across rotations of a central start platform such that the concordance between head and world reference frames was systematically disrupted. We hypothesised that this manipulation would allow us to unambiguously identify spatial reference frames during active sound localisation.

### 3.1 Simulation analysis

In order to confirm our expectations about how the firing of world and head centered neurons would be altered by sound position and start platform rotation we first simulated responses using model head and world centered units (see methods). For each unit, we computed the spatial tuning expected according to the location of sounds in the world (“world frame”, Fig. 2a) or the angle of the sound relative to the head (“Head frame”); by design, each unit was strongly modulated only in its native frame. We visualised neural firing across all possible head and world positions (Fig. 2b). In such plots the neural responses recorded to sound locations for any given platform location form a diagonal slice through the space of head and world positions, and recording at different platform locations allows the full 2D space to be quantified. Within this space the head-centered unit elicits a horizontal band of activity, illustrating invariance to the position of the sound in the world and a dependence only on the location of the sound angle relative to the head. In contrast, world-centered coding appears as a vertical band, indicating selectivity for world angle and invariance to head angle. Fig. 2c quantifies these patterns using model fit (Eq. 4) and model preference (Eq. 5). For a head-centered unit, model fit of head is close to 1 and model fit of world is close to 0, with model preference values near +1, whereas for a world-centered unit the opposite pattern is observed, with model preference values near −1.

### 3.2 Summary of neural data

We recorded 1,777 units from two ferrets (F1901 and F1905), of which 596 (33.5%) were stimulus driven. By animal and hemisphere (Table 1), F1901 yielded 413 units in the left hemisphere (166 stimulus driven; 40.2%) and 337 units in the right hemisphere (123 stimulus driven; 36.5%), whereas F1905 yielded 439 units in the left hemisphere (156 stimulus driven; 35.5%) and 588 units in the right hemisphere (151 stimulus driven; 25.7%).

**Table 1.** Summary of units per animal and hemisphere.

| Animal | Hemisphere | Total units (#) | Driven units (#) |
| --- | --- | --- | --- |
| F1901 | Left | 413 | 166 |
| F1901 | Right | 337 | 123 |
| F1905 | Left | 439 | 156 |
| F1905 | Right | 588 | 151 |

### 3.5 Neural representations across reference frames

Following the simulations, we applied the same analysis to neural recordings (Fig. 3). Visualisation revealed examples of firing rate patterns that were consistent with both head-centered (Fig. 3a-c) and world-centred (Fig. 3d,e) predictions. Model preference values (Eq 5), reported in Fig.3g, which confirmed the frame assignment for each example, as well as the presence of mixed selectivity. (Fig 3.f).

### 3.4 Most units show head-centered tuning, but a consistent minority show world-centered tuning

Across the two animals and four hemispheres, we found that the majority of driven units were to spatially modulated, with 9% of driven neurons being untuned. All tuned units were further classified as head centered, world-centered, or mixed, as shown in Fig. 4. In F1901 (Fig. 4a), 289 of 750 units were stimulus driven; of these, 208 (72%) were head centred, 48 (16.6%) were world centred, and 33 (11.4%) were not tuned. In F1905 Fig. 4b, 307 of 1,027 units were stimulus driven; of these, 232 (75.6%) were head centred, 49 (16.0%) were world centred, and 26 (8.5%) were not tuned. Hemisphere-wise counts were consistent with these totals (F1901 right hemisphere: 83/123 head, 25/123 world, 15/123 not tuned; F1901 left: 125/166 head, 23/166 world, 18/166 not tuned. F1905 right: 113/151 head, 33/151 world, 5/151 not tuned; F1905 left: 119/156 head, 16/156 world, 21/156 not tuned).

To quantify reference-frame preference, we plotted model fit (%) for the head model against the world model for all tuned units and summarised these distributions per ferret (Fig. 4c-d). Units were classified as strong head centred (Model Preference > +0.2), strong world centred (Model Preference

< -0.2) or mixed (|Model Preference| ≤ 0.2). In F1901 (Fig. 4c), of 256 tuned units, 187 (73%) were strong head centred, 29 (11.3%) were strong world centred and 40 (15.6%) were ambiguous. In F1905 (Fig. 4d), of 281 tuned units, 219 (77.9%) were strong head centred, 34 (12.1%) were strong world centred and 28 (10.0%) were ambiguous. The combined scatter plots show a dense cluster below the identity line (head dominant), a smaller cluster above it (world dominant) and a band along the diagonal (ambiguous).

### 3.5 Temporal dynamics of driven neurons

As outlined in Section 2.3.5, after screening for stimulus responsiveness and assessing spatial tuning in 150 ms windows, we classified units according to their temporal response properties (Fig. 4e-f). Units were assigned to four categories: onset only (responsive at onset but not offset), offset only (responsive at offset but not onset), both (responsive in both phases) and mixed across sessions. The mixed category denotes units that met the across-session criterion only when onset and offset sessions were considered together (for example, responsive at onset in one session and at offset in another) but did not meet the “at least two sessions” requirement within either phase alone. All categories were retained for subsequent analyses. In total, F1901 (Fig. 4e) yielded 192 onset-only, 20 offset-only, 51 both and 26 mixed across session units; F1905 (Fig. 4f) yielded 214 onset-only, 19 offset-only, 58 both and 16 mixed across session units.

We then analysed units classified as having both onset and offset responses to determine whether units’ spatial reference frame was stable over response epochs. As shown in Fig. 5a heat map, reference frame identity was largely preserved across phases; most units remained head-centred (53 units), with only one-unit remained world-centred.

**Figure 5.**
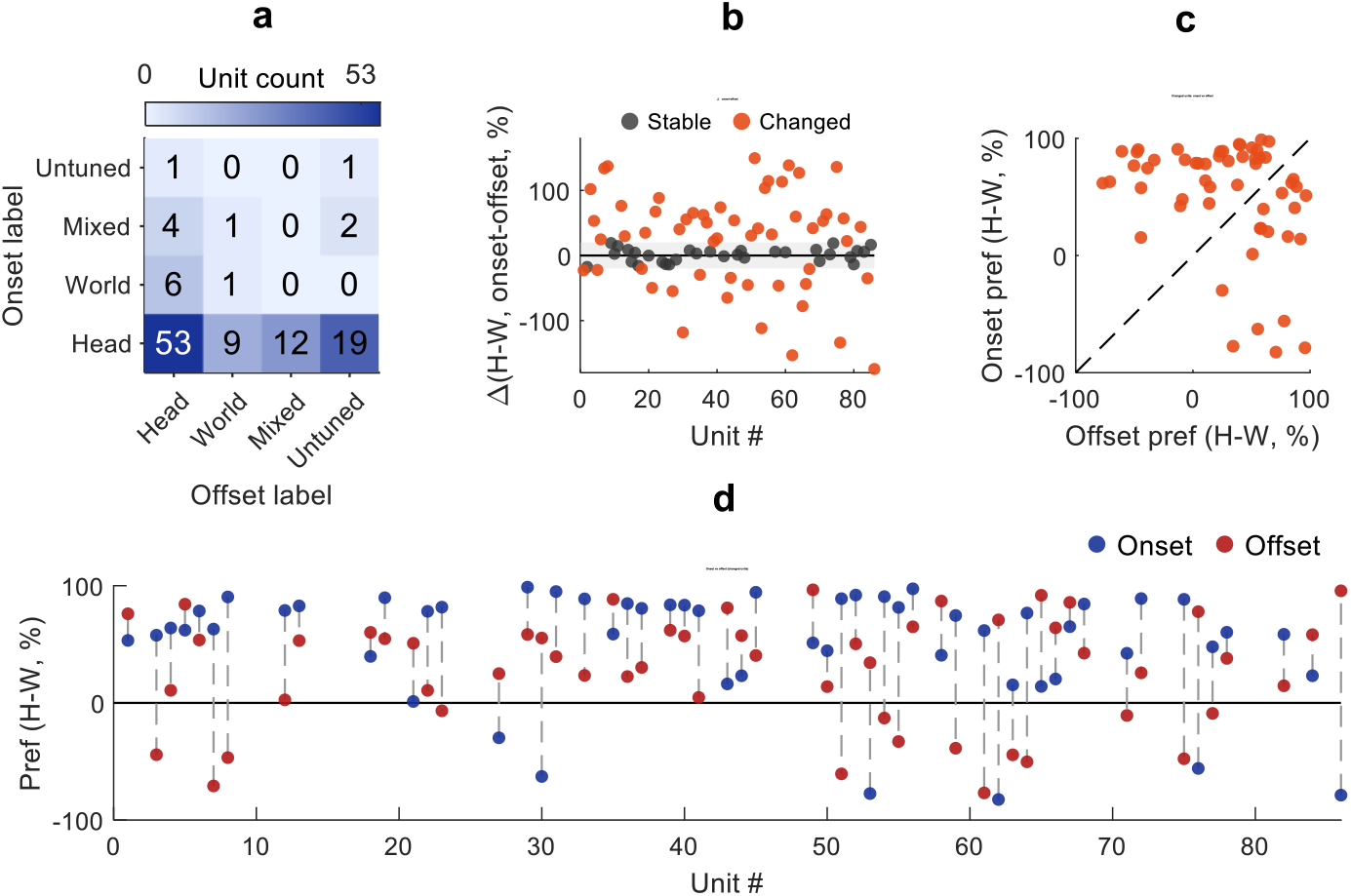
Onset-offset stability of reference-frame coding in “both-phase” units (both ferrets combined). (a) transition heat maps showing counts of onset-offset categorical transitions (Head, World, Mixed (ambiguous), untuned). (b) Change in continuous head-versus-world model preference (onset minus offset) for each tuned unit; dark grey points, stable units (|Δ| ≤ 20 percentage points); orange points, units with |Δ| > 20. (c) Onset versus offset model preference for changed units only (orange), plotted relative to the unity line (dashed). (d) Paired onset (blue) and offset (red) preferences for changed units arranged by unit number.

Having quantified temporal changes at the categorical level, we then imposed a more stringent, model-based criterion. For each unit that was tuned in both windows, we calculated the change in head-versus-world model preference between onset and offset (onset minus offset; Fig. 5b). Units with an absolute change of ≤20 percentage points were classified as stable, and those with a change exceeding 20 percentage points were classified as changed. This metric captures both units that transition between categorical labels and units that retain the same label but exhibit a marked reduction in absolute preference, such as Head-labelled units at both onset and offset that show a substantial loss of head-centred dominance. Using this criterion, 38 units in F1901 met the inclusion criteria, with 16/38 (42.1%) classified as stable and 22/38 (57.9%) showing a detectable change. In F1905, 48 units passed the same criteria, with 15/48 (31.2%) classified as stable and 33/48 (68.8%) as changed. Pooled across both animals, 31/86 units (36.0%) were stable, whereas 55/86 units (64.0%) exhibited a shift in reference-frame preference between onset and offset responses.

In Fig. 5c, plotting onset versus offset preference for the changed units only shows that most points fall above the unity line, indicating that head-versus-world preference is typically stronger at onset than at offset. Fig. 5d displays the same changed units as paired onset-offset values along the unit axis, illustrating that changes usually reflect a reduction in head-centred dominance over time to ambiguous, with a minority of units reversing the sign of their preference.

## 4. Discussion

In this short report we report the neural activity observed in auditory cortex in two animals performing a head-centered sound localisation task. In keeping with previous work ^15, 16^, we found a dominance of neurons encoding sound location in head centered coordinates, but a consistent minority that represented space in a world centered reference frame. Animals were required to discriminate source location (defined relative to the animals’ head) across rotations of the central start platform. By pseudorandomly varying the starting orientation across testing sessions we were able to break the relationship between sound source position relative to the animals’ head, and the speaker location in the world. World centered units showed tuning to specific locations within the arena, independently of the starting location of the animal.

This work replicates our previous findings and extends them to a more controlled setup where we were able to sample a full range of head and world positions. Thus, in contrast to our previous work, where animals showed a bias in the direction that they faced within the testing arena we were able to obtain a largely unbiased selection of head positions. Our previous study in freely moving, passively listening animals reported a strikingly similar proportion of world-centered neurons (∼15% of spatially modulated neurons, which were roughly 50% of driven neurons). In this study a larger proportion of driven responses were spatially tuned (80-90%) but the proportion of world-centered neurons was largely similar, despite the animal being required to disregard information about where the sound source was in the world to effectively perform the task. In this study our separation of head and world-centered tuning was less obviously bimodal than in ^16^, the mixed representations warrant further investigations as to whether these responses reflect weaker spatial modulation or joint coding of spatial and task relevant variables.

We were able to record neural activity across both hemispheres. Analysis of neural responses by hemisphere provided no evidence of hemispheric specialisation with the proportion of tuned neurons and the division of tuned neurons into different reference frames being very similar. In this task animals were required to discriminate front from rear sound sources; a situation in which neurons are particularly dependent on spectral cues, as sounds in the midline elicit identical binaural cues. Repeating these experiments with animals making left right decisions would be interesting, in order to compare the spatial tuning functions and spatial reference frames of recorded neurons.

There are some limitations to our current work: because we were only able to rotate the starting platform between sessions, this work required spike sorting neural responses across several weeks of data. While our modelling accounted for baseline changes in firing rate (that may be real, or an artefact of electrode drift / gliosis) by using session as a random effect, this may lead us to underestimate any effects of head-direction tuning that may result in “mixed” representations.

Our analyses are restricted to a head-centred behavioural task, which likely underestimates the contribution of world-centered neural tuning. We hypothesise that extending this approach to animals trained in a world-centered sound localisation task may reveal a greater proportion of world-centered responses and allow us to further explore whether spatial reference frames evolve over time. In these experiments we did not manipulate visual or vestibular inputs, so the mechanistic origin of the late, more world centered component remains unresolved.

## Acknowledgments

This work was funded by:

Wellcome Trust and Royal Society award 098418/Z/12/A.

Wellcome Career Development award 227480/Z/23/Z. European Research Council, 771550 SOUNDSCENE

Biotechnology and Biological Sciences Research Council, BB/N001818/1.

We are grateful for the excellent technical support provided by the staff at the Royal Veterinary College.

